# Integrating multimodal data through interpretable heterogeneous ensembles

**DOI:** 10.1101/2020.05.29.123497

**Authors:** Yan Chak Li, Linhua Wang, Jeffrey N. Law, T. M. Murali, Gaurav Pandey

## Abstract

**Motivation:** Integrating multimodal data represents an effective approach to predicting biomedical characteristics, such as protein functions and disease outcomes. However, existing data integration approaches do not sufficiently address the heterogeneous semantics of multimodal data. In particular, early and intermediate approaches that rely on a uniform integrated representation reinforce the consensus among the modalities, but may lose exclusive local information. The alternative late integration approach that can address this challenge has not been systematically studied for biomedical problems.

**Results:** We propose Ensemble Integration (EI) as a novel systematic implementation of the late integration approach. EI infers local predictive models from the individual data modalities using appropriate algorithms, and uses effective heterogeneous ensemble algorithms to integrate these local models into a global predictive model. We also propose a novel interpretation method for EI models. We tested EI on the problems of predicting protein function from multimodal STRING data, and mortality due to COVID-19 from multimodal data in electronic health records. We found that EI accomplished its goal of producing significantly more accurate predictions than each individual modality. It also performed better than several established early integration methods for each of these problems. The interpretation of a representative EI model for COVID-19 mortality prediction identified several disease-relevant features, such as laboratory test (blood urea nitrogen (BUN) and calcium) and vital sign measurements (minimum oxygen saturation) and demographics (age). These results demonstrated the effectiveness of the EI framework for biomedical data integration and predictive modeling.

**Availability:** Code and data are available at https://github.com/GauravPandeyLab/ensemble_integration.

## 1 Introduction

*Multimodal data* are a collection of diverse types of data that capture complementary aspects of a biomedical entity and its characteristics of interest (Boehm *et al*. (2021)). For instance, a protein may be characterized by its amino acid sequence, three-dimensional structure, evolutionary history and interactions with other proteins. These different types of information may be used to infer the function(s) of the protein (Pandey *et al*. (2006)). Similarly, Electronic Health Record (EHR) systems contain diverse types of clinical data, such as from questionnaires, imaging and laboratory tests, that collectively provide a comprehensive view of a patient’s health (Jensen *et al*. (2012)). These multimodal data are expected to be complementary. For instance, data from questionnaires provides baseline information of a patient’s health, while laboratory test measurements indicate their current health state. Due to this complementarity, integrating these multimodal data can result in a more comprehensive understanding and accurate predictions of biomedical characteristics or outcomes, such as the functions of proteins and disease phenotypes of patients (Boehm *et al*. (2021); Krassowski *et al*. (2020); Hasin *et al*. (2017)).

However, the integration and predictive modeling of multimodal data are still challenging because of the heterogeneity of the individual data modalities, which are usually structured differently and have different semantics (Zitnik *et al*. (2019); Gligorijević and Pržulj (2015)). For instance, gene expression data can be structured as matrices, each entry of which individually denotes the expression level of a gene in a sample. In contrast, string-formatted amino acid sequences are unstructured, since each position in these sequences needs to be studied in the context of its surrounding positions to infer its biological role. Several data types also present their own challenges, such as the high dimensionality of gene expression profiles, which can make their analysis difficult (Saeys *et al*. (2007)). Similar issues exist with the multimodal data in EHR systems as well. Due to the varying semantics and challenges of the individual modalities, it has been challenging to identify a uniformly effective prediction method for diverse multimodal data (Zitnik *et al*. (2019)).

Several data integration methods have been proposed to address the challenge of heterogeneity among multimodal data (Zitnik *et al*. (2019); Gligorijević and Pržulj (2015)). These methods can be generally categorized into three groups, namely *early, intermediate* and *late* integration (Fig. 1). Early integration strategies (Fig. 1a) first combine the multimodal data into a uniform intermediate representation, such as a network (Caldera *et al*. (2017); Wang *et al*. (2014)), which is then used for prediction and other analysis purposes. Intermediate strategies (Fig. 1b) jointly model the multiple datasets and their elements, such as genes, proteins, etc., also through a uniform intermediate representation. These uniform representations reinforce the consensus among modalities, but may obfuscate the local signal exclusive to each individual modality, and thus adversely affect prediction performance (Greene and Cunningham (2009); Libbrecht and Noble (2015)).

**Fig. 1:**
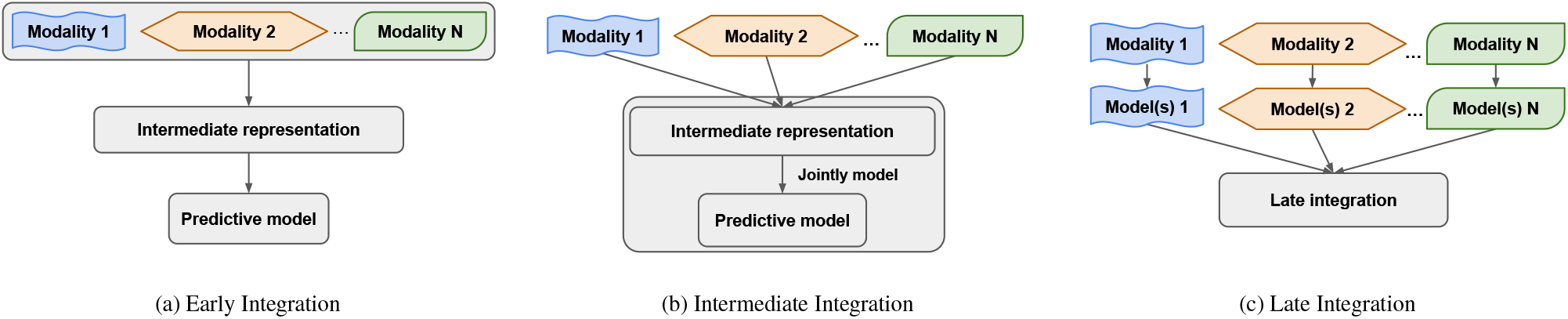
Overview of early, intermediate and late approaches for integrating multimodal data. (a) The early approach combines multiple datasets into an intermediate representation, from which predictive models can be inferred. (b) The intermediate approach jointly models the multiple datasets and their elements through an intermediate representation. (c) The late approach first builds local model(s) for each individual modality, which are then integrated into the final predictive model.

In view of these limitations of the early and intermediate approaches, the late approach (Fig. 1c) offers an alternative path to improved prediction performance by first deriving specialized local predictive models from the individual modalities, and aggregating those models (Zitnik *et al*. (2019); Gligorijević and Pržulj (2015)). This approach has the potential to capture maximal information in the individual modalities into the local predictors, whose aggregation then builds consensus, thus effectively utilizing both the commonalities and diversity among the modalities. However, this approach has not been systematically implemented and studied for biomedical data integration and predictive modeling problems (Zitnik *et al*. (2019)).

In this work, we propose *heterogeneous ensembles* as a novel systematic realization of the late integration strategy to build effective integrative predictive models from multimodal biomedical data. These ensembles can aggregate an unrestricted number and types of local models, thus providing an effective framework for aggregating diverse information captured in these models (Whalen *et al*. (2016)). Significantly, this approach differs from homogeneous ensembles, such as random forest (Breiman (2001)) and boosting (Schapire and Freund (2013)), which typically aggregate only one type of individual models, e.g., decision trees in random forests. Furthermore, homogeneous methods learn individual models as a part of the ensemble process, and thus cannot integrate models that have been derived independently *a priori*. The advantages of heterogeneous ensembles have been demonstrated in several biomedical applications, such as protein function prediction (Whalen *et al*. (2016); Wang *et al*. (2018); Stanescu and Pandey (2017)), enhancing the predictive power of DREAM Challenges (Sieberts *et al*. (2016, 2021)) and modeling infectious disease epidemics (Ray and Reich (2018)). However, these applications were limited to individual (unimodal) datasets.

In the current work, we leverage the flexibility of the heterogeneous ensemble framework to advance predictive modeling from *multimodal data* in an approach that we name *Ensemble Integration* (EI; Fig. 2). Specifically, EI uses the same heterogeneous ensemble methods as used for unimodal data to integrate local models derived from individual data types. Through this process, EI is capable of incorporating both the consensus and diversity among the individual modalities.

**Fig. 2:**
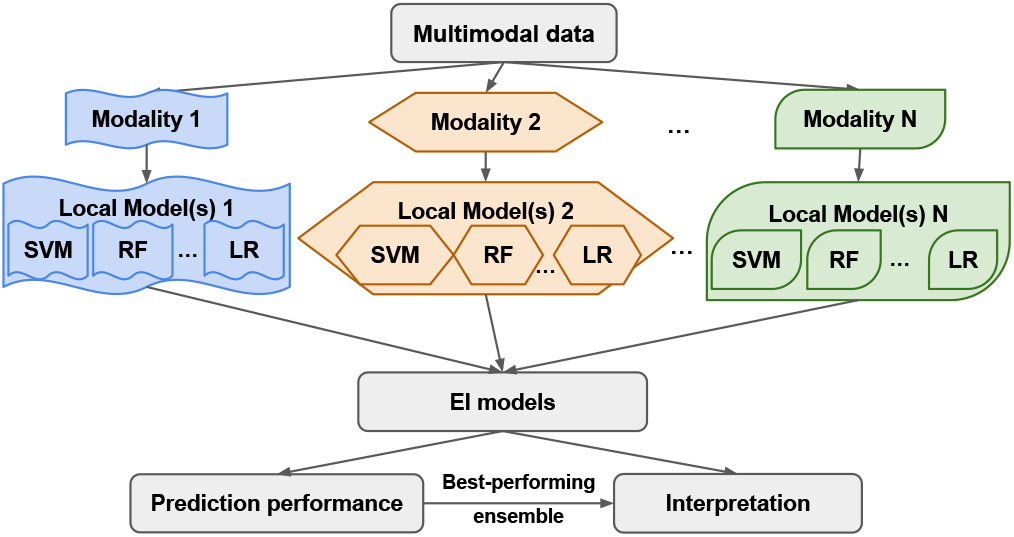
Overview of the Ensemble Integration (EI) framework for multimodal data. In the implementation of EI tested in this work, we used ten standard binary classification algorithms, such as support vector machine (SVM), random forest (RF) and logistic regression (LR), as implemented in Weka (Frank *et al*. (2005)), to derive sets of local predictive models 1, 2,…,N from the data modalities 1, 2,…,N. We then applied the stacking and ensemble selection methods to these local models to generate the EI models. These models generated prediction scores for the entities and multimodal data of interest that were evaluated to assess their performance. Finally, we used our novel interpretation method to identify the features that contributed the most substantially to the best-performing EI model’s predictions.

A challenge associated with multimodal data integration using EI is that the ensembles generated may be complex and difficult to interpret, not unlike other sophisticated machine learning-based models (Doshi-Velez and Kim (2017)). The interpretation of these ensembles is important for understanding the rationale behind their predictions and generating trust for the user. To address this challenge, we also propose a novel *interpretation framework* for EI-based models built from multimodal data.

We tested the prediction and interpretation capabilities of EI on two diverse and challenging problems: protein function prediction (PFP) and disease outcome (COVID-19 mortality) prediction. Specifically, we evaluated EI’s performance at predicting the functions (GO terms) of human proteins from multimodal datasets in the STRING database (Szklarczyk *et al*. (2021)). We also tested the effectiveness of EI for predicting the likelihood of a COVID-19 patient’s mortality (death) from the disease using the multimodal data collected in a treating hospital’s EHRs (Wynants *et al*. (2020)). We compared EI’s performance with those of established data integration methods that have been used for these problems. Finally, in addition to evaluating EI’s prediction abilities, we also used our ensemble interpretation method to reveal the key features that contributed to the EI-based predictive model of COVID-19 mortality, and verified the relevance of these features from the literature.

## 2 Methods

Here, we describe the methodologies and datasets used in our development and evaluation of the EI framework and other approaches. We followed the Data, Optimization, Model and Evaluation (DOME) recommendations (Walsh *et al*. (2021)), as detailed in the opening subsection of Supplementary Materials.

### 2.1 Ensemble Integration (EI)

Fig. 2 shows the general EI framework for multimodal data, as well as the implementation of EI built and tested in this work. First, predictive models were trained on each data modality; we refer to these as *local models* below. We used ten established binary classification algorithms implemented in Weka (Frank *et al*. (2005)) for training local models. These algorithms were AdaBoost, Decision Tree (J48), Gradient Boosting, K-nearest Neighbors, Support Vector Machine (SVM), Random Forest, Logistic Regression (LR), Rule-based Classification (PART), Naive Bayes and Voted Perceptron. All the algorithms were used with their default parameters in Weka, with the exception of specifying C=0.001 for SVM and M=100 for LR to control time to convergence. To handle potential imbalance between the classes being predicted, the entities belonging to the majority negative class in each modality were randomly under-sampled to balance the number of positive and negative class entities before training the local models. The corresponding test sets retained the same class ratio.

Next, using the base predictions generated by the local models, EI used the following heterogeneous ensemble methods to build the integrative predictive model.

- *Mean aggregation*, which calculates the ensemble output as the mean of the base prediction scores.
- The iterative *ensemble selection* method of Caruana *et al*. (2004, 2006) (*CES*), which starts with an empty set as the current ensemble. In each iteration, CES adds the local model that improves the current ensemble’s prediction performance by the largest amount. The iterative process, which samples the local models with replacement, continues until there is no further gain in performance.
- *Stacking* (Sesmero *et al*. (2015)), which uses the base predictions as features to train a second-level predictive model (meta-predictor) as the final ensemble. We used eight established binary classification algorithms available in Python’s sklearn library (Pedregosa *et al*. (2011)) to train meta-predictors. These algorithms were AdaBoost, Decision Tree, Gradient Boosting, K-nearest Neighbors, SVM, Random Forest, LR and Naive Bayes. We also included the XGBoost classifier (Chen and Guestrin (2016)) as a meta-prediction algorithm. All the algorithms were executed using their default parameter settings, with the exception of the linear kernel being used with SVM to control the runtime.

Thus, each execution of EI yielded eleven models, i.e., one each based on mean aggregation and CES, and nine based on stacking. These EI models then generated the final prediction scores for the entities of interest. These scores were evaluated to assess the models’ performance.

### 2.2 Interpretation of EI models

To aid the interpretation of EI models and build trust in them, we propose a novel method to identify the key features in the various data modalities that contribute the most to the model’s predictions (Supplementary Fig. 1). This method first quantifies the contributions the features in the individual modalities (local features) make to the corresponding local models. It then quantifies the contribution of each local model to the EI ensemble. Finally, it combines these contributions to determine the most important features for the EI model.

Algorithmically, the method calculates the *local feature ranks (LFRs)* from the local models and the *local model ranks (LMRs)* from the EI model (Algorithm 1) as follows. The *LFR*s for each local model are calculated based on the difference of the performance of the model when each feature is held out. This calculation is carried out using the ClassifierAttributeEval function in Weka. The calculation of *LMR*s depends on the type of ensemble algorithm used to build the EI model under consideration. If the model under consideration is based on mean aggregation, all the local models are assigned the same *LMR*s. If it is based on CES, the local models are assigned *LMR*s based on how many times they are included in the final ensemble. Finally, if the EI ensemble is based on stacking, *LMR*s are determined using the permutation importance of each constituent local model (Breiman (2001)). This importance measure is calculated as the average change in performance of the ensemble when the local model’s output is randomly permuted a hundred times and as many ensembles are retrained. This calculation is conducted using the permutation_importance function in the sklearn library (Pedregosa *et al*. (2011)). *LMR*s are then determined based on the descending order of the permutation importance values. Due to the varying number of local models and features these ranks are calculated for, all the ranks are normalized into percentile ranks that range from 1*/n* to 1, where *n* is the total number of local models in the ensemble or the total number of features in the modality being considered.

#### Algorithm 1: Calculate percentile ranks of local models in an ensemble

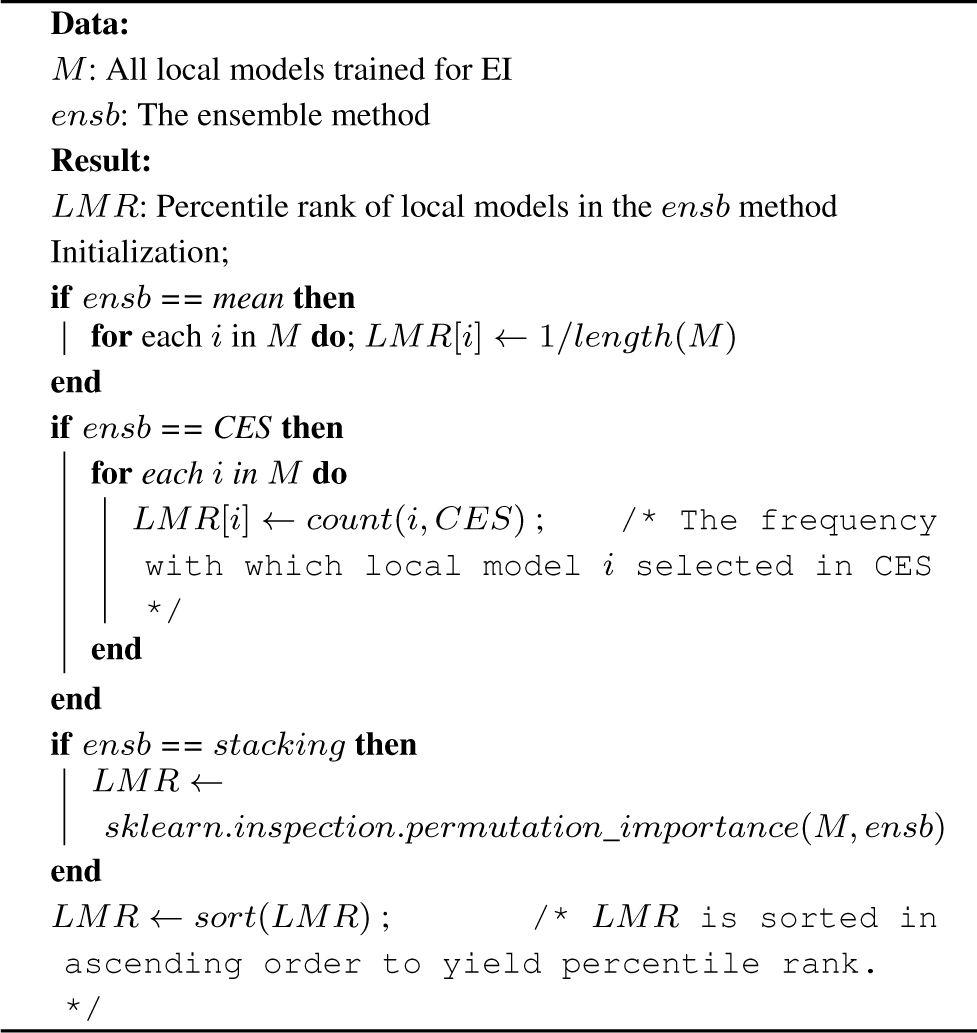

#### Algorithm 2: Calculate final ranking of features for an ensemble

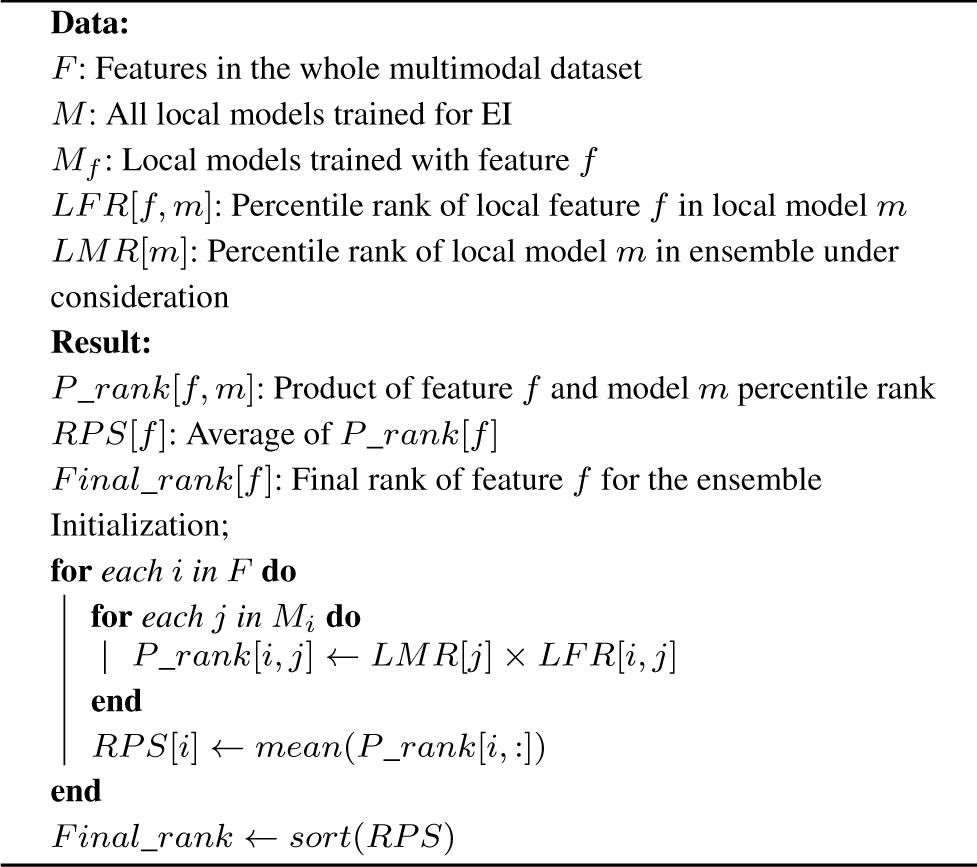

Next, we combined these two lists of ranks to compute the feature ranking for the EI model as shown in Algorithm 2. We first calculated the product of the *LMR* and *LFR* for each valid pair of local model and feature. We then averaged these products for each local feature across all the local models to generate its rank product scores (*RPS*s), which quantified the feature’s overall contribution to the EI model. All the features in all the modalities were then sorted in ascending order of *RPS*s to determine the final ranking of the features.

Note that this interpretation method is applicable to any EI model. However, in order to focus its use in our experiments, we applied it only to the best-performing ensemble algorithm within EI for the label of interest (Fig. 2). This ensemble algorithm was used to train an EI model on the whole dataset, which was then interpreted.

### 2.3 Biomedical problems used for evaluating EI

To evaluate the effectiveness of EI for biomedical problems, we tested the framework on two representative problems, one each from bioinformatics and clinical informatics. These problems were protein function prediction from STRING data (Szklarczyk *et al*. (2021)), and COVID-19 mortality prediction from EHR data, respectively. Below, we provide details of these problems, as well as the associated multimodal datasets used.

#### 2.3.1 Protein function prediction from STRING data

As discussed in the Introduction, a variety of complementary data modalities can be used to predict the functions of proteins (Pandey *et al*. (2006)), and the most accurate predictions are often obtained by integrating these modalities (Zitnik *et al*. (2019)). Thus, protein function prediction (PFP) is an ideal problem to test EI’s effectiveness for integrating multimodal data and producing accurate predictions. We followed the most commonly used definition of protein functions as Gene Ontology (GO) terms (Radivojac *et al*. (2013)). We predicted annotations of human proteins to these terms using diverse multimodal datasets from version 11.5 of the STRING database (Szklarczyk *et al*. (2021)) released in August 2021. These datasets consist of pairwise functional associations between thousands of proteins from several species, including human. These associations are derived from the multi-omic data sources listed in Table 1. Since these data sources provide complementary information about protein function (Pandey *et al*. (2006); Zitnik *et al*. (2019); Radivojac *et al*. (2013)), the associations derived from them in STRING are also multi-modal.

**Table 1:** Modalities of STRING data used for protein function prediction. For each modality of STRING, the description and number of features are included in the table.

| Modality | # features | Description |
| --- | --- | --- |
| PPI | 17,901 | Protein-protein interactions |
| Curated database | 10,501 | Interactions in curated databases |
| Co-expression | 18,618 | Co-expression relationships between proteins |
| Neighborhood | 3,974 | Co-location in the same genomic neighborhood |
| Co-occurrence | 2,936 | Co-occurrence of orthologs in other genomes |
| Fusion | 6,053 | Fusion of orthologs in other genomes |

We tested the effectiveness of EI for predicting protein function (GO term annotations) from these multimodal STRING datasets. For all the individual datasets, the adjacency vector of each protein was used as its feature vector for PFP model training and evaluation, since this representation has been shown to be effective for automated network-based gene/protein classification (Liu *et al*. (2020)). If a certain protein was not included in any of the STRING datasets, its corresponding feature vector was assigned to be all zeros before integration and prediction. EI was executed for each GO term individually on the multimodal datasets.

We compared EI’s performance with those of established network-based early integration algorithms, namely Mashup (Cho *et al*. (2016)) and deepNF (Gligorijević *et al*. (2018)) following the methodology described in Supplementary Fig. 2. Mashup and deepNF use graph propagation- and deep learning-based algorithms, respectively, to derive an integrated network corresponding to the input networks. In our study, these algorithms were applied to the STRING data described above to generated corresponding integrated networks. The same adjacency vector as feature vector representation was used for the integrated networks generated using these methods. This representation yielded feature vectors of lengths 800 and 1200 from the Mashup and deepNF integrated networks. Note that these networks already represent a layer of integration of the STRING data. Thus, to generate predictive models of GO term annotations from these networks that were consistent with the single layer of integration in EI (Supplementary Fig. 2), we used the local modeling algorithms listed in Section 2.1. Finally, to assess the value of integrating multimodal data, we also predicted annotations from each of the STRING datasets individually using the same heterogeneous ensemble algorithms as included in EI (Section 2.1). Again, these heterogeneous ensembles represented a single layer of integration of the base models derived from the individual modalities (Supplementary Fig. 2).

EI and the baseline methods were used to predict the annotations of 18,866 human proteins to 2,139 GO Molecular Function and Biological Process terms that were used to annotate at least fifty human proteins in the data released on January 13th, 2022. This lower limit was chosen to reduce the variability and enhance the reliability of the results obtained. For each GO term, the proteins annotated to it with non-IEA evidence codes were defined as positive examples. We defined negative examples as follows: a protein *p* was labeled as a negative example for a term *t* in an ontology *o* (e.g., Molecular Function) if *p* had at least one annotation to a term in *o*, but was not manually annotated (i.e., with a non-IEA evidence code) to *t* nor to its ancestors or descendants (Mostafavi *et al*. (2008)). For each term, only the positively or negatively annotated proteins were used for the evaluation of EI and the baseline methods.

It is noteworthy that GO terms are organized within hierarchical ontologies (The Gene Ontology Consortium (2017); Ashburner *et al*. (2000)), with deeper terms representing more specific functions and annotating fewer proteins. These properties of a GO term, namely its depth and the number of proteins annotated with it, have been shown to substantially influence the performance of PFP methods (Zhou *et al*. (2019)). Thus, to compare the performance of EI and the baseline methods more comprehensively, we also assessed how their performance varied with these properties. For this, we first compared the performances of the prediction methods across GO terms grouped by the number of human genes annotated to them. In our consideration set, there were 166 terms with more than 1000 annotations, 162 with 500 to 1000 annotations, 388 with 200 to 500 annotations, 544 with 100 to 200 annotations and 879 with 50 to 100 annotations. Since these numbers of proteins annotated to GO terms varied over several orders of magnitude, we also considered a more normalized measure of these number known as information content of a term (Zhou *et al*. (2019)). For a term *t*, this quantity is defined as *-* log_10_(*p*(*t*)), where *p*(*t*) is the probability of a human protein being annotated with *t*. Finally, the depth of GO term *t* was defined as the shortest path from the root of the corresponding ontology to *t*. The depth and information content of GO terms were calculated using the GOATOOLS package (version 1.0.3) (Klopfenstein *et al*. (2018)).

#### 2.3.2 COVID-19 mortality prediction from EHR data

The COVID-19 pandemic has infected over 570 million individuals and caused over 6 million deaths globally, as of July 22*nd*, 2022, as per the Johns Hopkins University Coronavirus Resource Center (Dong *et al*. (2020)). It thus represents the most serious public health threat the world has faced in a long time. For a patient being treated for this disease, it is immensely useful for healthcare providers to have an estimate of the patient’s outcomes, since they can adapt the treatment accordingly (Wynants *et al*. (2020)). The multimodal data collected from these patients into their EHRs, such as data collected at admission, co-morbidity information, vital signs and laboratory test measurements, represent useful sources of information that can be integrated and used for predicting these outcomes.

We tested EI’s ability to address this need. Specifically, we used EI to predict the likelihood of an individual dying from COVID-19 (*mortality*), the most serious outcome, for patients treated at Mount Sinai between March 15 and October 1, 2020 from the data modalities in the EHR system described in Table 2 (details of the features are in Supplementary Table 1). These modalities and features were selected and processed by expert clinicians and informaticians, and provided by the Mount Sinai Data Warehouse. Since vital signs and laboratory tests were measured multiple times for each patient, we used their respective first values recorded during the first 36 hours of hospitalization to enable early outcome prediction, as recommended by another study (Vaid *et al*. (2020)). Only features with fewer than 30% missing values were included in the analysis. Any remaining missing values in each modality were imputed using the KNNImpute method (Troyanskaya *et al*. (2001)) with K set to 5. The categorical features in all the modalities were transformed to numerical values by one-hot encoding. The continuous features were normalized into z-scores. The resultant number of features included in each modality are specified in the descriptions above. Using the above data, we tested EI for predicting if a patient died from COVID-19 (mortality) during the course of their hospitalization. This outcome was determined using the value of the Boolean “deceased” flag in the patient’s EHR. In our dataset covering 4, 783 patients, 1, 325 (27.7%) passed away from the disease.

**Table 2:** Modalities of electronic health record (EHR) data used for predicting mortality due to COVID-19. For each modality of EHR data, the description and number of features are included in the table.

| Modality | # features | Description |
| --- | --- | --- |
| Admission | 23 | Baseline information collected from patients at hospital admission, including age, race/ethnicity, sex, and some vital signs, such as oxygen saturation and body temperature. |
| Co-morbidities | 22 | Simultaneous presence of other common conditions in patients, such as asthma and obesity, that can affect COVID-19 trajectory and outcomes. |
| Vital signs | 9 | Clinically appropriate maximum and/or minimum values of heart rate, body temperature, respiratory rate, diastolic and systolic blood pressure, and oxygen saturation level that define a patient’s clinical state during hospitalization. |
| Laboratory tests | 44 | Values of laboratory tests, such as the number of white blood cells and platelets in blood, conducted on samples taken from patients during the course their hospitalization to monitor their internal health. |

Specifically, we compared EI’s performance with that of an early integration approach proposed by another study (Vaid *et al*. (2020)). This approach concatenated the feature vectors from the individual modalities for a given patient, and built mortality prediction models using XGBoost (Chen and Guestrin (2016)), which is considered the most effective method for tabular data (Shwartz-Ziv and Armon (2021)). The default parameters specified in the Python xgboost library (Chen and Guestrin (2016)) were used for this baseline method. Additionally, similar to the PFP experiments, we also considered heterogeneous ensembles built on the individual data modalities as alternative baselines.

Finally, we used the novel method described in Section 2.2 to identify the features in the above data modalities that contributed the most to the EI predictive models for COVID-19 mortality. We also compared these features with those that contributed the most to the XGBoost model for the same outcome. For this, as in Vaid *et al*. (2020)’s study, we calculated the mean absolute SHAP value (Lundberg *et al*. (2020)) for all the features in the XGBoost model, and ranked them in descending order of this value to identify the most important features. We then conducted the Fisher’s exact test to calculate the statistical significance of the overlap between the two feature tests.

### 2.4 Evaluation methodology

All the heterogeneous ensembles, both from EI and baseline approaches, were trained and evaluated in a five-fold nested cross-validation (Nested CV) setup (Whalen *et al*. (2016)) in the above experiments. In this setup, the whole dataset is split into five outer folds, which are further divided into inner folds. The inner folds are used for training the local models, while the outer folds are used for training and evaluating the ensembles. Nested CV helps reduce overfitting during heterogeneous ensemble learning by separating the set of examples on which the local and ensemble models are trained and evaluated (Whalen *et al*. (2016)). The Mashup and deepNF baselines in PFP, and XGBoost baseline in the COVID-19 mortality prediction experiments were executed in the standard 5-fold cross-validation setup.

All the GO terms and COVID-19 mortality outcomes predicted in our experiments were unbalanced, as is common knowledge for PFP (Radivojac *et al*. (2013); Zhou *et al*. (2019)). To assess the performance of the prediction methods for the more challenging minority positive class in each experiment, we used *F*_*max*_ in all our evaluations. *F*_*max*_ is the maximum value of F-measure across all prediction score thresholds among the combinations of precision and recall for the positive class, and has been recommended for PFP assessment by the CAFA initiative (Radivojac *et al*. (2013); Zhou *et al*. (2019)). We also we measured the prediction performances for both problems in terms of the precision and recall yielding the *F*_*max*_ value reported. Furthermore, since our PFP experiments involved evaluations on over 2,000 GO terms, we also statistically compared the performances of EI and the baselines using Friedman and Nemenyi tests (Demšar (2006)) to assess the overall performance of all the methods tested.

Finally, as explained in the EI interpretation method (Section 2.2), performance assessment was also needed for calculating *LMR*s in EI ensembles, as well as for calculating *LFR*s for features included in the local models. For both these calculations, we used the Area Under the Precision-Recall Curve (AUPRC), since it was the only class imbalance-aware performance measure available as an option in the sklearn and Weka functions used (Section 2.2).

The code implementing all the above methods and the data used are available at https://github.com/GauravPandeyLab/ensemble_integration.

## 3 Results

Below, we describe and analyze the results obtained in our experiments on protein function and COVID-19 mortality prediction.

### 3.1 Protein function prediction

Fig. 3 shows the distributions of the *F*_*max*_ scores of all the protein function prediction (PFP) approaches tested in Section 2.3.1. For each approach, this figure shows a box plot denoting the distribution of one *F*_*max*_ score for each GO term. This score is for the algorithm implementing the approach that performed the best for that GO term, e.g., stacking using logistic regression for Ensemble Integration for GO:0000976 (transcription cis-regulatory region binding). As explained in Section 2.3.1, each of the prediction approaches tested here, namely EI, deepNF and Mashup with local modeling algorithms and the individual STRING modalities with heterogeneous ensembles, represented a single layer of data integration (Supplementary Fig. 2). Thus, the selection of the algorithm that yielded the best performance for each approach for each GO term is intended to present the corresponding strongest integrative predictor for each term, and thus facilitate consistency across the diverse approaches evaluated.

**Fig. 3:**
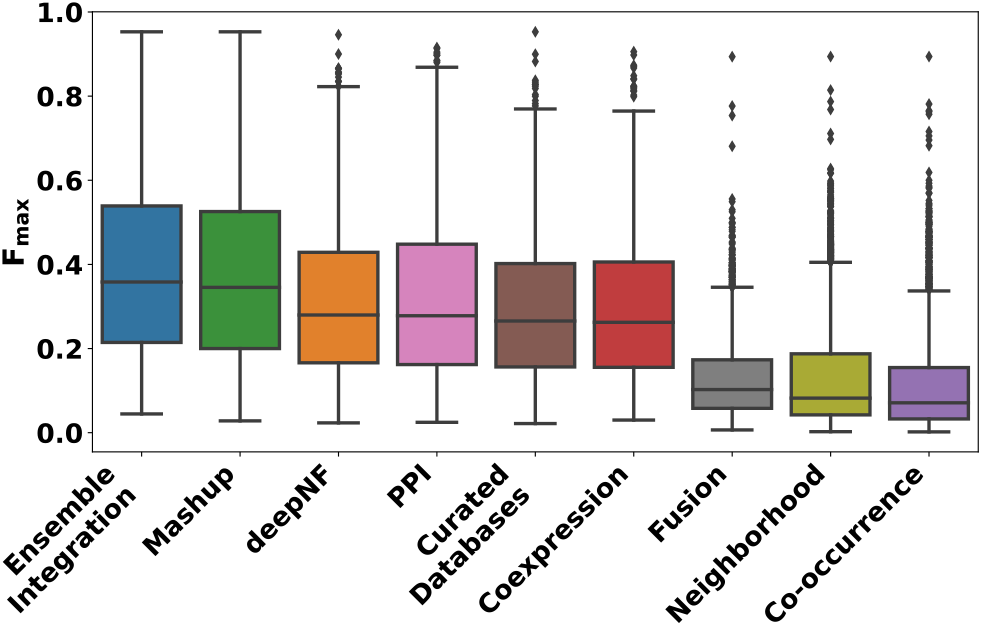
The distributions of the performances of the protein function prediction approaches tested in this work. This performance was measured in terms of the *F*_*max*_ score for each of the 2,139 GO terms. For EI and the individual modalities, the score for the best performing heterogeneous ensemble method for each GO term is shown here. For the Mashup and deepNF early integration methods, the score of the best local modeling algorithm for each term is shown.

These results show that the performance of Ensemble Integration (EI) was significantly better than those of Mashup and deepNF (Friedman-Nemenyi FDR = 9.34 × 10^*-*14^ and *<* 2 × 10^*-*16^, respectively), as well as each of the individual STRING data modalities (Friedman-Nemenyi FDR *<* 2 × 10^*-*16^). These results were generally consistent with those obtained in terms of the precision and recall yielding the above *F*_*max*_ values (Supplementary Fig. 4).

We also examined how the performance of EI, deepNF and Mashup varied with the depth (Fig. 4a) and information content (Fig. 4b) of the GO terms included in our evaluation. Across all depths and information content levels, EI consistently performed better than both deepNF and Mashup, exemplified by the higher median *F*_*max*_ values across all the GO term subsets. We also assessed the performance of all the prediction approaches across GO term subsets grouped by the number of human genes annotated to them (Supplementary Fig. 3). Although the performance of all the approaches deteriorated, as expected, for terms with fewer annotations, EI generally performed well, and better than the other approaches. The only exception to this observation was the set of terms with the fewest annotations (50–100), where Mashup produced significantly more accurate predictions than EI.

**Fig. 4:**
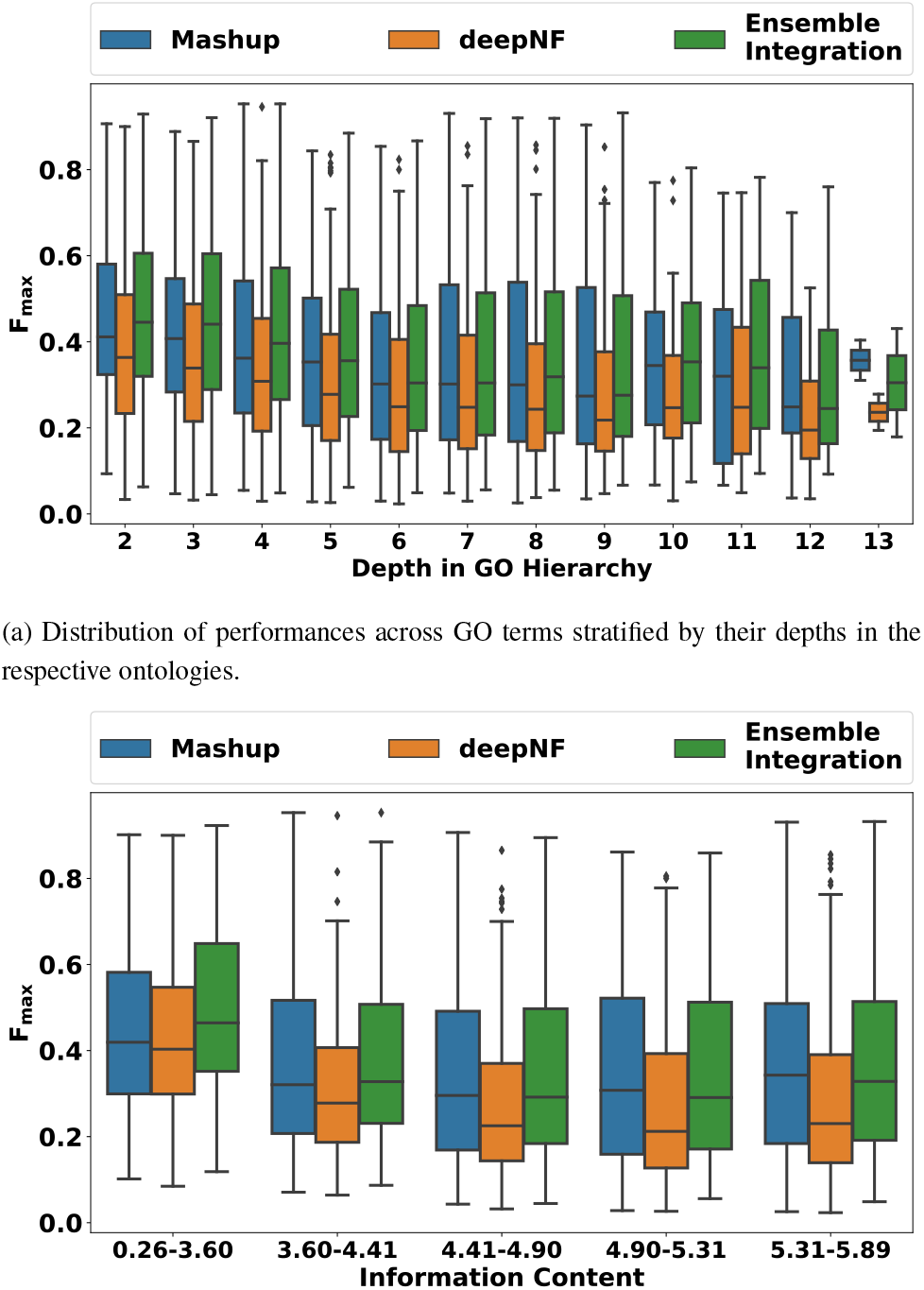
Distribution of the performances (*F*_*max*_ scores) of Ensemble Integration (EI), deepNF and Mashup for GO terms at varying depths and information content levels. The depth and information content values of the 2,139 GO terms included in this experiment were calculated using the GOATOOLS package (version 1.0.3) (Klopfenstein *et al*. (2018)).

Collectively, these results indicated that EI was able to achieve the goals of data integration and predictive modeling for PFP more effectively than other alternate approaches.

Finally, to gain insight into which ensemble methods performed the best for EI, we analyzed this distribution for all the GO terms tested (Supplementary Fig. 5). Stacking with random forest and logistic regression were generally the best-performing EI methods, consistent with observations in our previous work with single datasets (Whalen *et al*. (2016); Wang *et al*. (2018)).

### 3.2 COVID-19 mortality prediction

Fig. 5 shows the distributions of the performance of the various implementations of EI and heterogeneous ensembles derived from the individual EHR data modalities for predicting the mortality over hospitalization outcome. Note that, unlike Fig. 3, the performance of all the implementations of each of the approaches are shown in the corresponding box plots, since these results are only for one outcome. Also shown is the performance of the baseline XGBoost method (dotted red line). As in the PFP experiments, EI performed significantly better than the individual modalities (Wilcoxon rank-sum FDR *<* 0.0082). Two of the eleven EI ensembles tested performed better than XGBoost. The best-performing EI method (stacking using logistic regression) scored a 0.011 (1.66%) higher *F*_*max*_ than XGBoost.

**Fig. 5:**
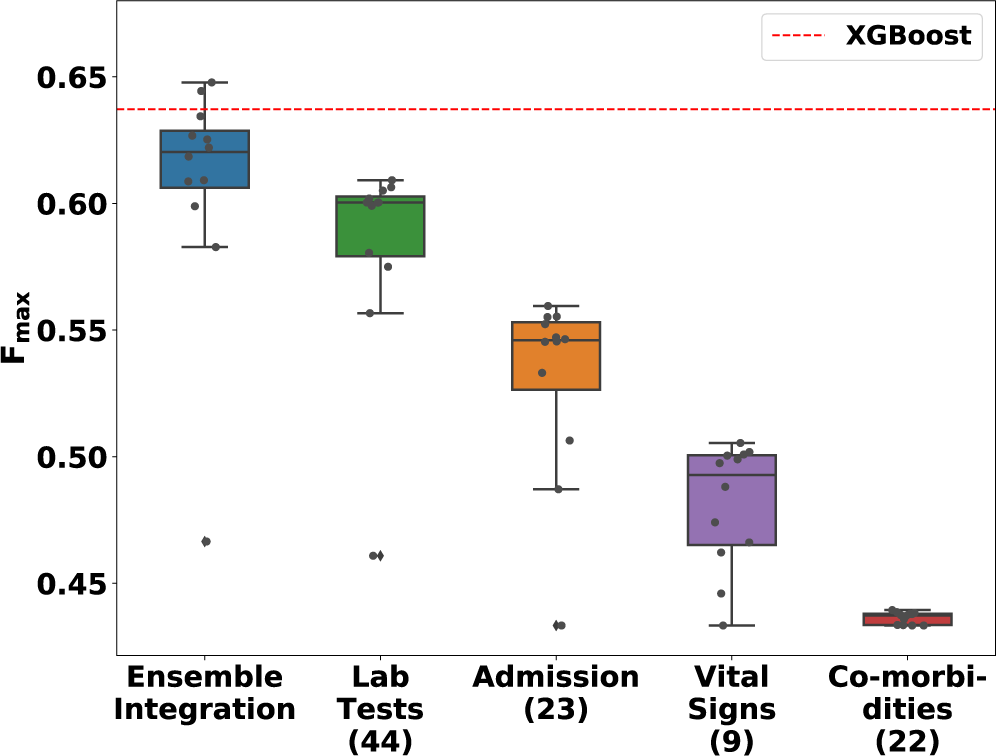
The distribution of *F*_*max*_ values of various ensemble models and XGBoost for predicting mortality from COVID-19 over a patient’s hospitalization. The performance distributions of Ensemble Integration (EI) and the heterogeneous ensembles derived from the individual EHR data modalities are shown as box-and-whisker plots. Each plot includes eleven ensemble models built using mean aggregation, CES, and nine stacking algorithms. The numbers in parentheses next to the names on the *x*-axis indicate the number of features in the corresponding modality. The dotted red line indicates the performance of the XGBoost model trained after concatenating the feature vectors in each individual modality.

We also examined the variation of precision and recall values for this best-performing EI method, the best-performing heterogeneous ensembles inferred from individual modalities and XGBoost (Fig. 6). The comparative results were generally consistent with those shown in Fig. 5, with EI generally performing well and slightly better than XGBoost. EI also achieved a slightly better balance between precision (0.569) and recall (0.752) at the point where F-measure was maximized than the other approaches, including XGBoost (precision=0.529, recall=0.801).

**Fig. 6:**
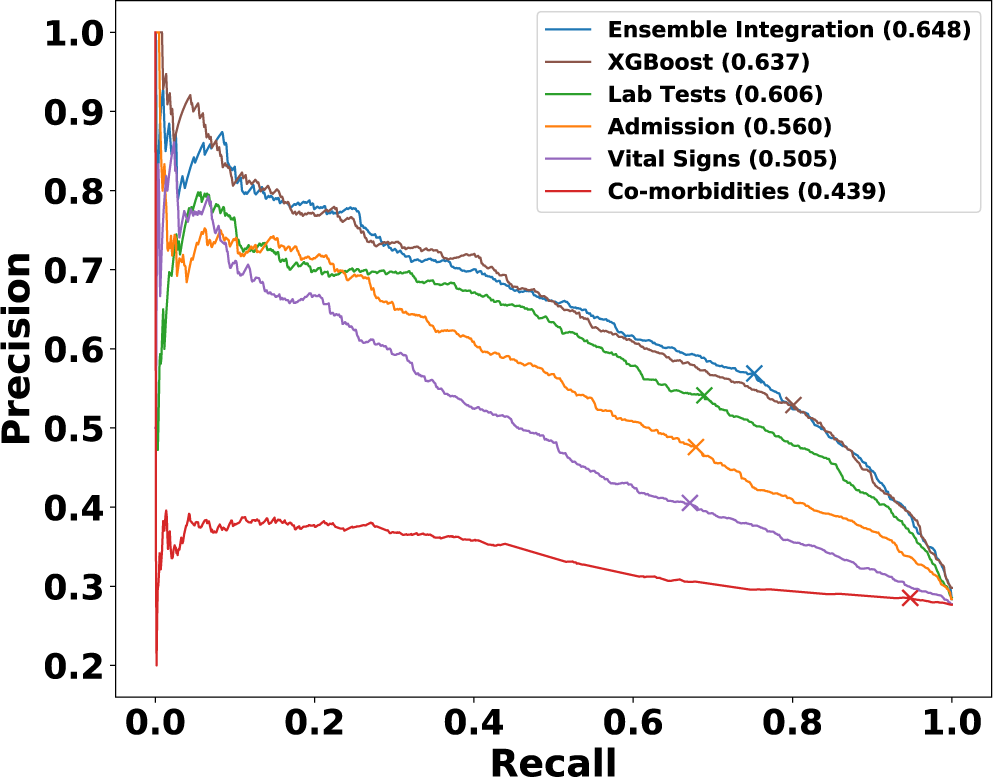
Precision-recall curves of representative models from EI, individual EHR modalities and XGBoost for predicting COVID-19 mortality. The values in parentheses show the *F*_*max*_ value of each of these models, and the cross marker on each curve indicates the precision and recall at which the corresponding *F*_*max*_ was obtained.

Overall, these results showed the slight advantage of EI over early integration for COVID-19 mortality prediction.

### 3.3 Interpretation of COVID-19 mortality prediction

Using the method described in Section 2.2, we also identified the ten features that contributed the most to the best-performing EI model for the COVID-19 mortality outcome (Table 3). These features included five from laboratory tests, three from admission, one from vital signs and one from co-morbidities. This distribution was consistent with the observation that that laboratory tests produced the most accurate predictions across all the modalities (Fig. 5).

**Table 3:**
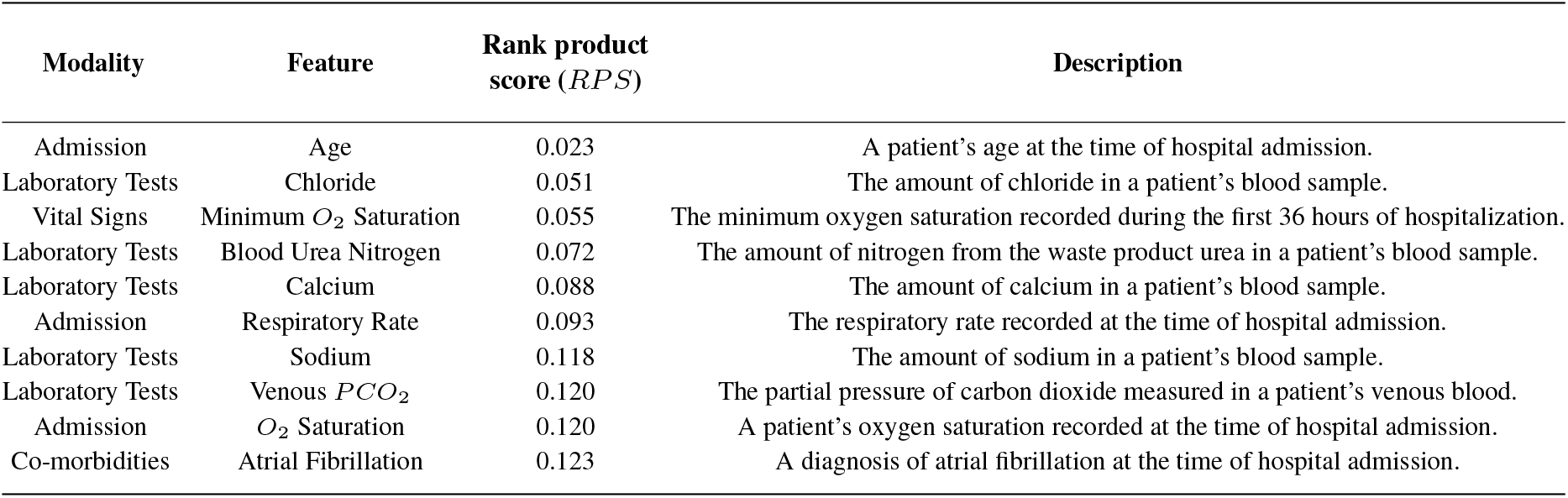
Ten highest contribution features identified from the best-performing EI model for predicting mortality due to COVID-19. The table provides detailed information on the features, including their names, the modality they were included in, and their description. The features are sorted in terms of increasing rank product scores (third column), which is the EI interpretation metric our method calculates (Section 2.2.)

We also identified the ten most contributing features in the XGBoost model (Supplementary Table 2). Four of these features, namely age at admission, minimum oxygen saturation among vital signs, and blood urea nitrogen and calcium measurements among laboratory tests, overlapped with the ten most predictive features for the EI model. This overlap was statistically significant (Fisher’s exact test p=0.0089, odds ratio=8.74).

The results in the above subsections illustrate the utility of our EI framework for addressing biomedical prediction problems, as well as interpreting the predictive models generated by the framework.

## 4 Discussion

We proposed a novel framework named Ensemble Integration (EI) to perform data integration and predictive modeling on heterogeneous multimodal data. In contrast to the more commonly used early and intermediate data integration approaches, EI adopts the late integration approach. In EI, one or more local models are derived from the individual data modalities using appropriate algorithms. These local models are then integrated into a global predictive model using effective heterogeneous ensemble methods, such as stacking and ensemble selection. Thus, EI offers the flexibility of deriving individually effective local models from each of the data modalities, which addresses challenges related to the differing semantics of these modalities. The use of heterogeneous ensemble methods then enables the incorporation of both the consensus and diversity among the local models and individual data modalities.

We tested EI for the diverse problems of protein function prediction (PFP) from multimodal STRING datasets and predicting mortality due to COVID-19 from multimodal EHR datasets. In both these experiments, we compared EI’s performance with those obtained from the individual data modalities, as well as established early integration methods, specifically deepNF and Mashup for PFP, and XGBoost for COVID-19 mortality prediction. In all these experiments, EI performed significantly better than the individual data modalities, showing that it accomplishes its data integration goals successfully. EI also performed better than the early integration approaches, most likely due to its ability to aggregate the complementary information encapsulated in the local models, which may be lost in the uniform integrated representations used by early and intermediate strategies.

We also proposed a novel interpretation method for EI models that ranks the features in the individual modalities in terms of their contributions to the EI model under consideration. We tested this method on the representative EI model constructed for predicting the COVID-19 mortality outcome. We found that several of the ten most important features identified for mortality due to COVID-19 were laboratory test measurements (Table 3), which was consistent with the observation that this modality yielded the most accurate predictions among all the individual modalities (Fig. 5). In particular, we found that the measurements of blood urea nitrogen (BUN) and calcium in patients’ blood samples were key features. These findings were consistent with prior research on the relevance of these measurements to mortality due to COVID-19 (Basheer *et al*. (2021); Lippi *et al*. (2020)). Patients’ age and minimum oxygen saturation were also found to be important features, again consistent with prior evidence (Price-Haywood *et al*. (2020); Yadaw *et al*. (2020); Berenguer *et al*. (2020); Pun *et al*. (2021)). We also analyzed the consistency of our list of most predictive features with the ten features found to be most predictive of the same outcome using Vaid *et al*. (2020)’s XGBoost method that was considered the baseline in our prediction experiments (Section 3.2). We found that four features, namely age at admission, minimum oxygen saturation among vital signs, and calcium and BUN measurements among laboratory tests, were common to both lists. This overlap was statistically significant, indicating that EI can identify important features consistent with other prediction methodologies, as well as reveal novel relevant features. A deeper investigation of the predictive features we identified can shed further light on the pathophysiology of COVID-19 and mortality due to the disease.

In summary, this paper presented the novel EI framework, and evidence in support of its ability to address challenging biomedical prediction problems. We also discussed challenges with the current work, and avenues for future work. Our efforts represent the first step in the systematizing and expanding the use of the late integration approach for complex, multimodal biomedical data.

Our work also has some limitations, which offer avenues for future work. First, although EI is capable of integrating both structured and unstructured data modalities, as well as a variety of local models derived from these modalities, we only tested EI with structured datasets and standard classification algorithms used to derive local models from these datasets. In the future, it would be valuable to test EI with unstructured data and specialized local models as well, such as label propagation on network data (Cowen *et al*. (2017)), convolutional neural networks (CNNs) derived from the biomedical images (Shen *et al*. (2017)) and recurrent neural networks (RNNs) trained on sequential or time series data (Geraci *et al*. (2017)). Furthermore, in our experimental evaluations, we only compared EI to the more commonly used early data integration approaches, namely Mashup and deepNF for PFP and XGBoost for COVID-19 mortality prediction. These comparisons should be expanded to intermediate integration approaches as well. Finally, our PFP evaluations were based on multimodal functional associations derived from diverse types of omic data in the STRING database. It would be valuable to evaluate the ability of EI for predicting protein function directly from the raw omic data types. Such expanded applications and evaluations will enable a more comprehensive assessment of the capabilities of EI.

There are some limitations with the EI interpretation method and its application as well. First, since predictive model interpretation can be subjective, and hence, not scalable (Doshi-Velez and Kim (2017)), we only tested this method on the COVID-19 datasets and mortality outcome, not the considerably more numerous (2,139) and diverse GO terms considered in the PFP experiments. To interpret the EI models built for these terms, it would be useful to prioritize these terms based on their relevance to the biological topic of interest.

Furthermore, the implementation of the proposed feature ranking method was based on AUPRC, since this was the only class imbalance-aware measure available in the Weka and sklearn functions used for determining local ranks of features and models respectively. However, since the basic principles of these rankings are general, they can be implemented with other performance measures, such as *F*_*max*_, as well using custom code. Also, due to the use of percentile ranks in the interpretation method, it is slightly biased in favor of modalities with larger sets of features. Thus, in addition to the clinical relevance of laboratory test measurements to monitoring patients’ COVID-19 status, this bias also played a role in these features being the highest ranked for the mortality outcome. This limitation can potentially be addressed by considering normalized versions of model and feature ranks.

## Acknowledgements

We thank Akhil Vaid and Sharon Nirenberg for their technical advice on our COVID-19 data processing and experiments, and Andrew DePass and Jamie Bennett for testing the EI code. The study was enabled in part by computational resources provided by Scientific Computing at the Icahn School of Medicine at Mount Sinai, Oracle Cloud credits and related resources provided by the Oracle for Research program.

## Funding

This work was supported by NIH grant# R01HG011407-01A1.

## Notes

https://github.com/GauravPandeyLab/ensemble_integration

### Competing Interest Statement

The authors have declared no competing interest.

### Summary of Updates

This version of the manuscript has been revised to provide more details and clarifications of the methods and results.

## References

Ashburner, M. et al. (2000). Gene ontology: tool for the unification of biology. the gene ontology consortium. Nat. Genet., 25(1), 25–29.

Basheer, M. et al. (2021). Clinical predictors of mortality and critical illness in patients with covid-19 pneumonia. Metabolites, 11(10).

Berenguer, J. et al. (2020). Characteristics and predictors of death among 4035 consecutively hospitalized patients with covid-19 in spain. Clin Microbiol Infect, 26(11), 1525–1536.

Boehm, K. M. et al. (2021). Harnessing multimodal data integration to advance precision oncology. Nature reviews. Cancer.

Breiman, L. (2001). Random forests. Machine Learning, 45(1), 5–32.

Caldera, M. et al. (2017). Interactome-based approaches to human disease. Current Opinion in Systems Biology, 3, 88–94.

Caruana, R. et al. (2004). Ensemble selection from libraries of models. Proceedings of the twenty-first international conference on Machine learning.

Caruana, R. et al. (2006). Getting the most out of ensemble selection. In Sixth International Conference on Data Mining (ICDM’06), pages 828–833.

Chen, T. and Guestrin, C. (2016). Xgboost: A scalable tree boosting system. In Proceedings of the 22nd ACM SIGKDD International Conference on Knowledge Discovery and Data Mining, KDD ‘16, pages 785–794, New York, NY, USA. Association for Computing Machinery.

Cho, H. et al. (2016). Compact integration of multi-network topology for functional analysis of genes. Cell Syst, 3(6), 540–548.

Cowen, L. et al. (2017). Network propagation: a universal amplifier of genetic associations. Nat Rev Genet, 18(9), 551–562.

Demšar, J. (2006). Statistical comparisons of classifiers over multiple data sets. Journal of Machine Learning Research, 7(1), 1–30.

Dong, E. et al. (2020). An interactive web-based dashboard to track COVID-19 in real time. The Lancet Infectious Diseases, 20(5), 533–534.

Doshi-Velez, F. and Kim, B. (2017). Towards a rigorous science of interpretable machine learning. arXiv: Machine Learning.

Frank, E. et al. (2005). Weka: A machine learning workbench for data mining., pages 1305–1314. Springer, Berlin.

Geraci, J. et al. (2017). Applying deep neural networks to unstructured text notes in electronic medical records for phenotyping youth depression. Evidence-Based Mental Health, 20(3), 83–87.

Gligorijević, V. and Pržulj, N. (2015). Methods for biological data integration: perspectives and challenges. J R Soc Interface, 12(112).

Gligorijević, V. et al. (2018). deepNF: deep network fusion for protein function prediction. Bioinformatics, 34(22), 3873–3881.

Greene, D. and Cunningham, P. (2009). A matrix factorization approach for integrating multiple data views. In W. Buntine, M. Grobelnik, D. Mladenić, and J. Shawe-Taylor, editors, Machine Learning and Knowledge Discovery in Databases, pages 423–438, Berlin, Heidelberg. Springer Berlin Heidelberg.

Hasin, Y. et al. (2017). Multi-omics approaches to disease. Genome Biology, 18(1), 83.

Jensen, P. B. et al. (2012). Mining electronic health records: towards better research applications and clinical care. Nat. Rev. Genet., 13(6), 395–405.

Klopfenstein, D. V. et al. (2018). Goatools: A python library for gene ontology analyses. Scientific Reports, 8(1), 10872.

Krassowski, M. et al. (2020). State of the field in multi-omics research: From computational needs to data mining and sharing. Frontiers in Genetics, 11, 1598.

Libbrecht, M. W. and Noble, W. S. (2015). Machine learning applications in genetics and genomics. Nat. Rev. Genet., 16(6), 321–332.

Lippi, G. et al. (2020). Electrolyte imbalances in patients with severe coronavirus disease 2019 (COVID-19). Ann Clin Biochem, 57(3), 262–265.

Liu, R. et al. (2020). Supervised learning is an accurate method for network-based gene classification. Bioinformatics, 36, 3457 –3465.

Lundberg, S. M. et al. (2020). From local explanations to global understanding with explainable ai for trees. Nature Machine Intelligence, 2(1), 56–67.

Mostafavi, S. et al. (2008). GeneMANIA: a real-time multiple association network integration algorithm for predicting gene function. Genome Biology, 9(Suppl 1), S4.

Pandey, G. et al. (2006). Computational Approaches for Protein Function Prediction: A Survey. Technical Report 06-028, University of Minnesota.

Pedregosa, F. et al. (2011). Scikit-learn: Machine learning in python. Journal of Machine Learning Research, 12(85), 2825–2830.

Price-Haywood, E. G. et al. (2020). Hospitalization and mortality among black patients and white patients with covid-19. N Engl J Med, 382(26), 2534–2543.

Pun, B. T. et al. (2021). Prevalence and risk factors for delirium in critically ill patients with COVID-19 (COVID-D): a multicentre cohort study. Lancet Respir Med, 9(3), 239–250.

Radivojac, P. et al. (2013). A large-scale evaluation of computational protein function prediction. Nature Methods, 10(3), 221–227.

Ray, E. L. and Reich, N. G. (2018). Prediction of infectious disease epidemics via weighted density ensembles. PLOS Computational Biology, 14(2), 1–23.

Saeys, Y. et al. (2007). A review of feature selection techniques in bioinformatics. Bioinformatics, 23(19), 2507–2517.

Schapire, R. E. and Freund, Y. (2013). Boosting: Foundations and algorithms. Kybernetes.

Sesmero, M. P. et al. (2015). Generating ensembles of heterogeneous classifiers using stacked generalization. WIREs Data Mining and Knowledge Discovery, 5(1), 21–34.

Shen, D. et al. (2017). Deep learning in medical image analysis. Annu Rev Biomed Eng, 19, 221–248.

Shwartz-Ziv, R. and Armon, A. (2021). Tabular data: Deep learning is not all you need.

Sieberts, S. K. et al. (2016). Crowdsourced assessment of common genetic contribution to predicting anti-tnf treatment response in rheumatoid arthritis. Nature Communications, 7(1), 12460.

Sieberts, S. K. et al. (2021). Developing better digital health measures of parkinson’s disease using free living data and a crowdsourced data analysis challenge. medRxiv.

Stanescu, A. and Pandey, G. (2017). Learning parsimonious ensembles for unbalanced computational genomics problems. Pac Symp Biocomput, 22, 288–299.

Szklarczyk, D. et al. (2021). The STRING database in 2021: customizable protein–protein networks, and functional characterization of user-uploaded gene/measurement sets. Nucleic Acids Research, 49(D1), D605–D612.

The Gene Ontology Consortium (2017). Expansion of the gene ontology knowledgebase and resources. Nucleic Acids Res., 45(D1), D331–D338.

Troyanskaya, O. et al. (2001). Missing value estimation methods for DNA microarrays. Bioinformatics, 17(6), 520–525.

Vaid, A. et al. (2020). Machine learning to predict mortality and critical events in a cohort of patients with covid-19 in new york city: Model development and validation. J Med Internet Res, 22(11), e24018.

Walsh, I. et al. (2021). DOME: recommendations for supervised machine learning validation in biology. Nature methods, 18(10), 1122–1127.

Wang, B. et al. (2014). Similarity network fusion for aggregating data types on a genomic scale. Nat Methods, 11(3), 333–337.

Wang, L. et al. (2018). Large-scale protein function prediction using heterogeneous ensembles. F1000Res, 7.

Whalen, S. et al. (2016). Predicting protein function and other biomedical characteristics with heterogeneous ensembles. Methods, 93, 92–102.

Wynants, L. et al. (2020). Prediction models for diagnosis and prognosis of covid-19: systematic review and critical appraisal. BMJ, 369.

Yadaw, A. S. et al. (2020). Clinical features of COVID-19 mortality: development and validation of a clinical prediction model. Lancet Digit Health, 2(10), e516–e525.

Zhou, N. et al. (2019). The CAFA challenge reports improved protein function prediction and new functional annotations for hundreds of genes through experimental screens. Genome Biology, 20(1).

Zitnik, M. et al. (2019). Machine learning for integrating data in biology and medicine: Principles, practice, and opportunities. Information Fusion, 50, 71–91.

