## Supplementary Material for "Integrating multimodal data through interpretable heterogeneous ensembles"

### Supplementary Materials

#### Compliance with DOME recommendations

Our study followed the Data, Optimization, Model and Evaluation (DOME) recommendations (Walsh *et al.* (2021)), as detailed below:

- **Data**: The evaluation of EI for protein function prediction (PFP) is based on publicly available STRING network and Gene Ontology annotation data, both described in Section 2.3.1. The same section also describes the number of proteins and features covered by the STRING data, as well as the distribution of proteins across the GO terms. We have shared all the data used in the PFP experiments with the public EI GitHub repository (https://github.com/GauravPandeyLab/ensemble_integration).

  The electronic health record (EHR) and outcome data used in the COVID-19 mortality prediction experiments were obtained from the Mount Sinai Data Warehouse, and were prepared by expert clinicians and informaticians. These data, including the distribution of patients over the values of the mortality outcome (alive and deceased) are decribed in Section 2.3.2. However, due to restrictions to protect patient privacy, we are unable to publicly share these data.
- **Optimization**: The algorithms used for building the local and EI ensemble models in this study are listed in Section 2.1. The default parameters of these algorithms in the respective public libraries they were adopted from (Weka and scikit-learn respectively) were used. The only exceptions were specifying C=0.001 for SVM and M=100 for LR to control time to convergence, based on our previous experience with these algorithms. However, we did not optimize the parameters of any of the prediction algorithms used for each dataset and/or label individually to avoid overfitting.

  All the training of the local and EI ensemble models was conducted in a nested cross-validation (Nested CV; Section 2.4) setup. In this setup, the whole dataset is split into five outer folds, which are further divided into inner folds. The inner folds are used for training the local models, while the outer folds are used for training and evaluating the ensembles. Nested CV also helps reduce overfitting during heterogeneous ensemble learning by separating the set of examples on which the local and ensemble models are trained and evaluated (Whalen *et al.* (2016)).

  All the algorithms and their parameters are included in the EI code provided at the public GitHub repository mentioned above. Users of the code are also able to change these settings as they desire.
- **Model**: Note that our study was focused on proposing and evaluating prediction algorithms, such as EI and benchmarks like deepNF and Mashup, and not to propose one or more specific models for our target problems. The only exception to this was the EI-based COVID-19 mortality prediction model that was interpreted in Section 3.3. We have shared this model through the GitHub repository mentioned above. We also hope that the results of the interpretation of this model will help shed light on COVID-19 pathophysiology, as well as help other researchers design and conduct related studies. More importantly, we hope that our EI framework provides a novel, reliable methodology for building specific models in other studies.
- **Evaluation**: As explained in Section 2.4, as well as relevant subsections of Section 3 (Results), we rigorously evaluated our proposed EI framework, and compared them with relevant benchmark approaches. Specifically, we used the Nested CV setup described above to fairly evaluation all the algorithms, as well as reduce overfitting in the process. We also used a variety of evaluation metrics, most prominently $F_{max}$, which was recommended by the Critical Assessment of Protein Function Annotation (CAFA) exercise (Radivojac *et al.* (2013)) for the evaluation of supervised methods for unbalanced classes, like in PFP. We also evaluated the consistency of our EI interpretation method with other methods and evidence in the literature (Section 3.3). Thus, consistent with the focus of our study, we rigorously evaluated all the algorithms tested, and assessed the results they generated.

We hope that the substantial details we have provided in accordance with the DOME recommendations for our study will aid its reproducibility and utility.

### Supplementary Figures

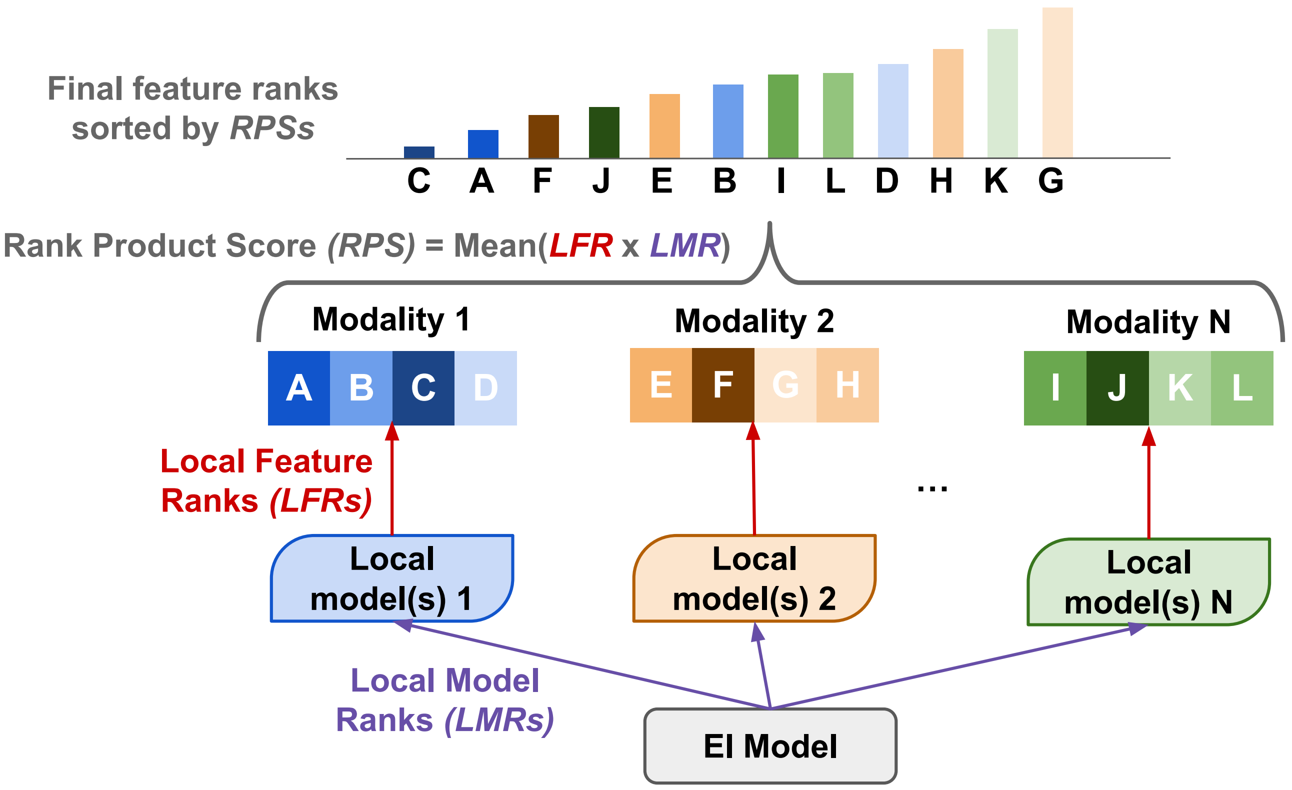


Supplementary Fig. 1: **Overview of the EI model interpretation method.** The method is based on local model ($LMR$s, purple arrow) and feature ($LFR$s, red arrow) ranks. $LMR$ denotes the importance of a local model derived from one of the data modality (e.g., Local model(s) 1 derived from Modality 1) to the final EI model, while $LFR$ denotes the contribution of each feature in the corresponding data modality (e.g., A-D in Modality 1) to a local model. The method averages the product of the $LMR$ and $LFR$ for each valid pair of local model and feature into a rank product score ($RPS$). The final ranking of all the features in terms of their importance is determined by sorting the $RPS$s in ascending order.

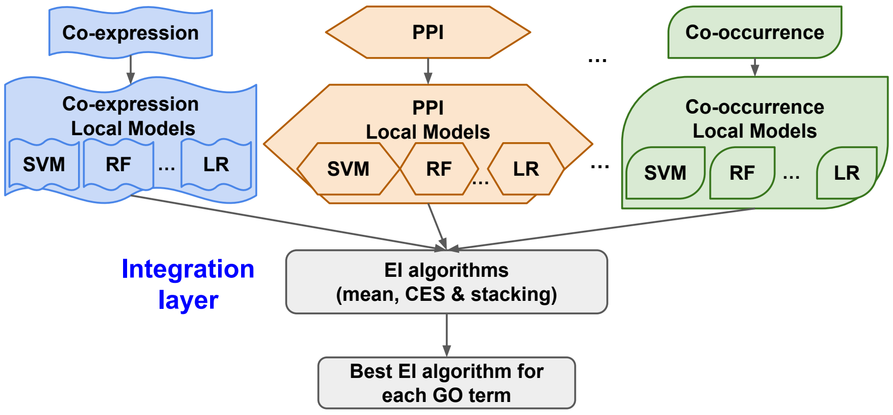


(a) Identifying the best-performing EI algorithm.

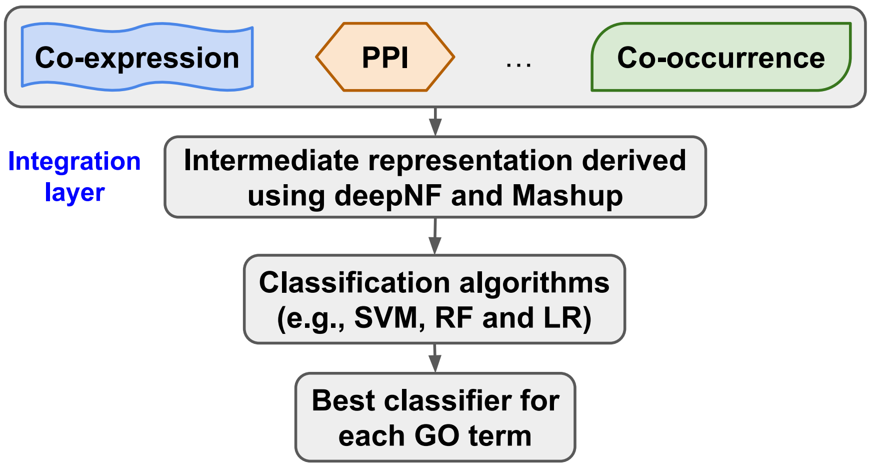


(b) Identifying the best-performing classification algorithm for the integrated networks (intermediate representations) derived using deepNF and Mashup.

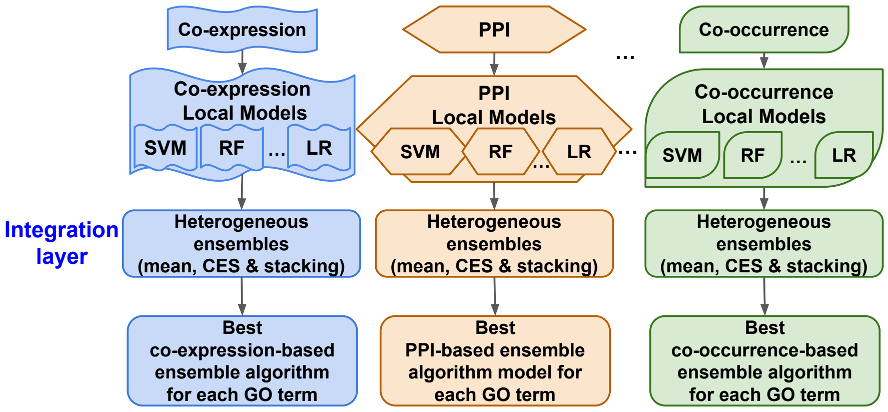


(c) Identifying the best-performing heterogeneous ensemble algorithms for the individual data modalities in STRING.

Supplementary Fig. 2: **Overview of the workflow for identifying the best-performing algorithms for protein function prediction.** These algorithms, namely (a) EI, (b) classifiers on integrated networks derived using deepNF and Mashup and (c) heterogeneous ensembles applied to the individual data modalities were applied to the STRING data as described in Section 2.3.1. Also marked in the workflows are the layers (steps) at which data and/or information were integrated. Based on the cross-validation results obtained, we identified and compared the best-performing algorithms in each of these categories for each GO term.

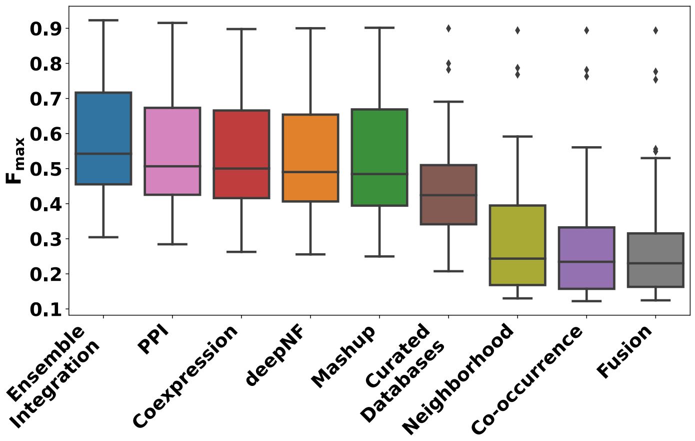


(a) GO terms with more than 1000 annotations (FDR of EI vs deepNF $= 9.05 \times 10^{-14}$, EI vs Mashup $< 2 \times 10^{-16}$, EI vs all individual modalities $< 3.19 \times 10^{-6}$).

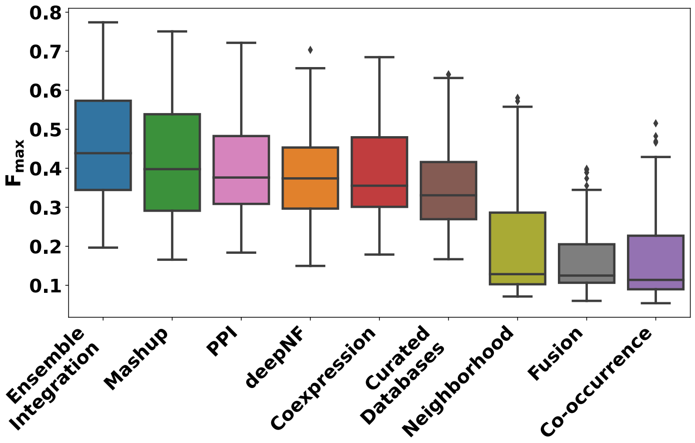


(b) GO terms with 500 to 1000 annotations (FDR of EI vs deepNF $= 8.86 \times 10^{-14}$, EI vs Mashup $= 1.09 \times 10^{-13}$, EI vs all individual modalities $< 8.77 \times 10^{-12}$).

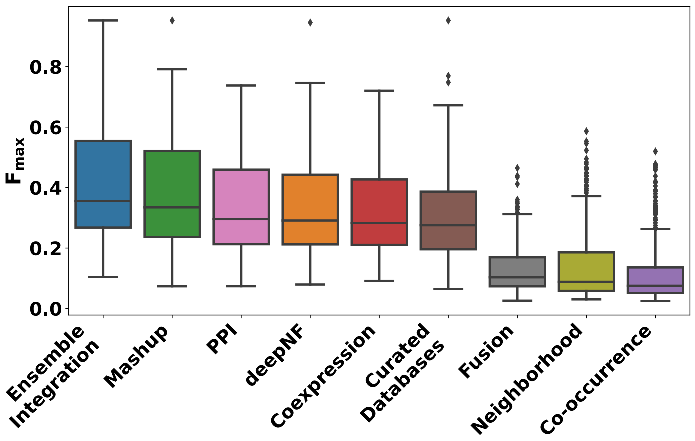


(c) GO terms with 200 to 500 annotations (FDR of EI vs deepNF $< 2 \times 10^{-16}$, EI vs Mashup $= 3.52 \times 10^{-10}$, EI vs all individual modalities: $< 2 \times 10^{-16}$).

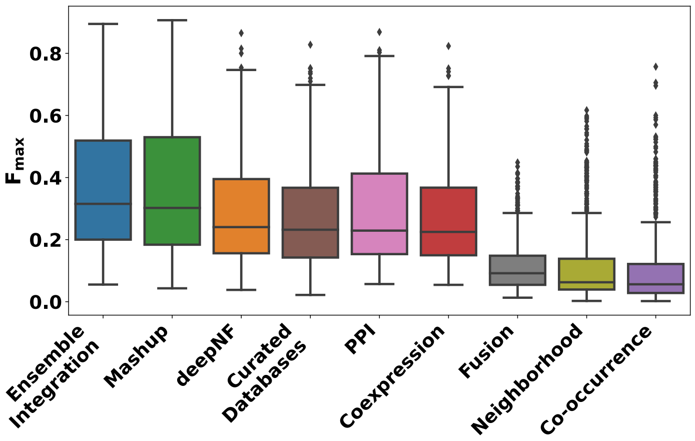


(d) GO terms with 100 to 200 annotations (FDR of EI vs deepNF $< 2 \times 10^{-16}$, EI vs Mashup $= 0.001$, EI vs all individual modalities $< 2 \times 10^{-16}$).

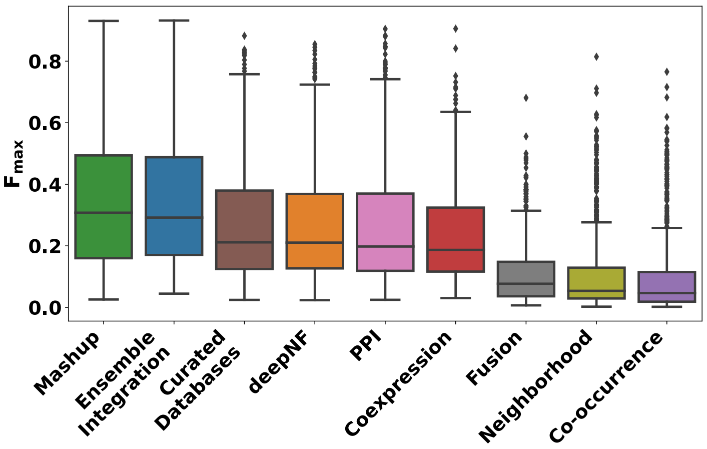


(e) GO terms with 50 to 100 annotations (FDR of EI vs deepNF $< 2 \times 10^{-16}$, EI vs Mashup $= 7.77 \times 10^{-4}$, EI vs all individual modalities $< 2 \times 10^{-16}$).

Supplementary Fig. 3: **Distributions of performances of the protein function prediction approaches tested in this work across GO terms grouped by the number of human genes annotated to them.** Performance was measured in terms of the $F_{max}$ score. Also shown are the FDR values representing the statistical significance of the comparative performance of EI vs deepNF, Mashup and individual STRING data modalities.

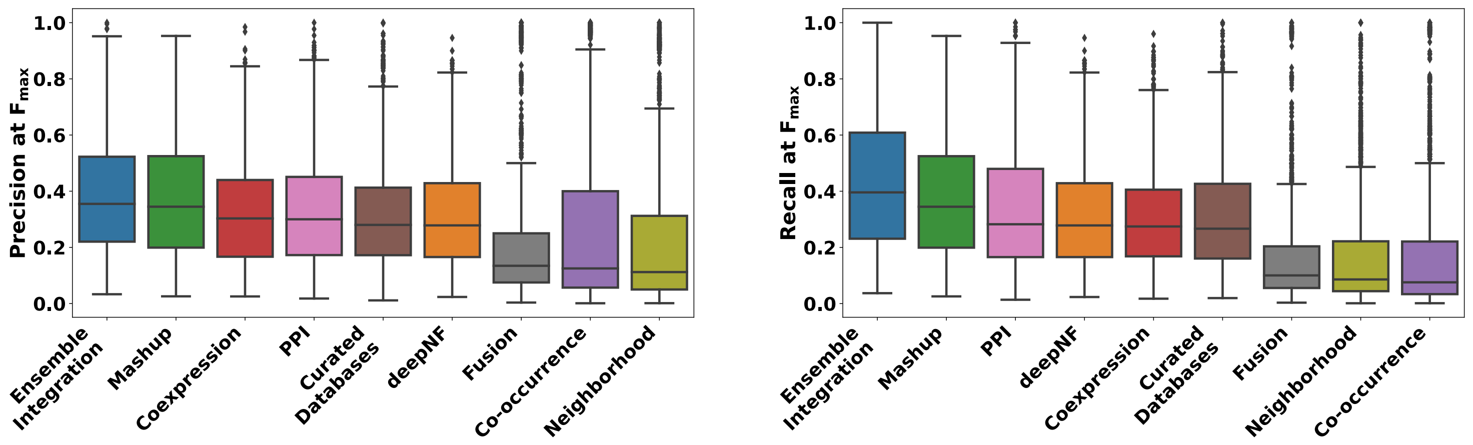


(a) Distribution of precision at $F_{max}$ across all the GO terms tested (FDR of EI vs deepNF $< 2 \times 10^{-16}$, EI vs Mashup $= 0.006$, EI vs all individual modalities $< 2 \times 10^{-16}$).

(b) Distribution of recall at $F_{max}$ across all the GO terms tested (FDR of EI vs deepNF $< 2 \times 10^{-16}$, EI vs Mashup $< 2 \times 10^{-16}$, EI vs all individual modalities $< 2 \times 10^{-16}$).

Supplementary Fig. 4: **Distributions of (a) precision and (b) recall yielding the $F_{max}$ values reported in Fig. 3 for all the protein function prediction approaches, data modalities and GO terms tested in this work.** Also shown are the FDR values representing the statistical significance of the comparative performance of EI vs deepNF, Mashup and individual STRING data modalities.

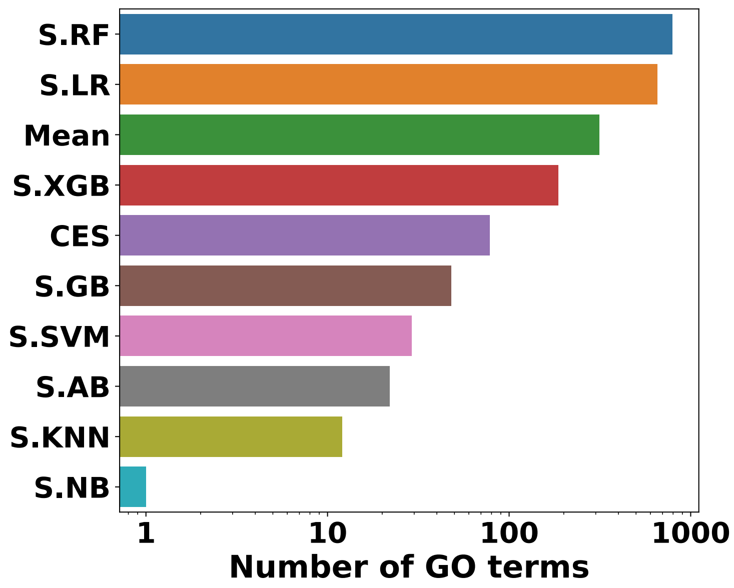


Supplementary Fig. 5: **Distribution of best-performing heterogeneous ensemble methods used within EI for protein function prediction.** This distribution was calculated across all the GO terms and the eleven ensemble methods tested. The Y-axis shows the names of the ensemble methods. The names with prefix ‘S.’ denote stacking with the classification algorithm named in the suffix, e.g., ‘S.RF’ stands for stacking with random forest. The X-axis shows the count (in logarithmic scale) of the GO terms for which each ensemble method showed the best performance. Note that stacking with decision tree (S.DT) is not shown here, since it was not found to be the best performer for any term, i.e., it’s count on the X-axis was zero.

**Supplementary Tables**

Table 1: Details of the clinical variables in electronic health records (EHRs), organized by the modalities they belonged to, that were used to predict mortality due to COVID-19 (Section 2.3.2 of the main text). Also provided are the units of the laboratory tests, as well as the exact or prefix of ICM-10-CM diagnosis code used for determining the values of the features in the co-morbidities modality.

**Admission (variables collected at the beginning of a patient's hospital encounter)**

| **Feature** | Description (units where applicable) |
|---|---|
| Age | The patient's age rounded down to nearest integer at the time of their hospital encounter. |
| Diastolic BP | The patient's first diastolic blood pressure reading taken during the encounter (mmHg). |
| Heart Rate | The patient's first recorded heart rate during the encounter (beats/minute). |
| Oxygen Saturation | The patient's first recorded oxygen saturation during the encounter (percentage). |
| Respiratory Rate | The patient's first recorded respiratory rate during the encounter (breaths/minute). |
| Systolic BP | The patient's first systolic blood pressure reading taken during the encounter (mmHg). |
| Temperature | The patient's first recorded temperature during the encounter (Fahrenheit). |
| Race/Ethnicity - Amercian Indian or Alaska Native | Binary variable indicating if the patient identified as Amercian Indian or Alaska Native. |
| Race/Ethnicity - Asian | Binary variable indicating if the patient identified as Asian. |
| Race/Ethnicity - Black or African-American | Binary variable indicating if the patient identified as Black or African-American. |
| Race/Ethnicity - White | Binary variable indicating if the patient identified as White. |
| Race/Ethnicity - Hispanic | Binary variable indicating if the patient identified as Hispanic. |
| Race/Ethnicity - Unknown | Binary variable indicating if the patient identified as unknown Race/Ethnicity. |
| Race/Ethnicity - Other | Binary variable indicating if the patient identified as other Race/Ethnicity. |
| Sex - Female | Binary variable indicating if the patient identified as female. |
| Sex - Male | Binary variable indicating if the patient identified as male. |
| Smoking Status - Never | Binary variable indicating if the patient never smoke. |
| Smoking Status - Not Asked | Binary variable indicating if the patient was not asked about their smoking status. |
| Smoking Status - Passive | Binary variable indicating if the patient's smoking status is passive. |
| Smoking Status - Quit | Binary variable indicating if the patient quit smoking. |
| Smoking Status - Yes | Binary variable indicating if the patient is still smoking. |

**Co-morbidities (binary variables indicating various morbidities diagnosed by ICD-10-CM codes)**

| **Feature** | **Description** |
|---|---|
| Acute Kidney Injury | Diagnosed by an ICD-10-CM code beginning with N17. |
| Acute Myocardial Infarction | Diagnosed by an ICD-10-CM code beginning with I21. |
| Acute Venous Thromboembolism | Diagnosed by an ICD-10-CM code beginning with I26 or I82.4. |
| Alcoholic/Non-alcoholic Liver Disease | Diagnosed by ICD-10-CM codes K75.81 or K76.0. |

(continued on the next page)

Table 1: (continued from the previous page)

| | |
|---|---|
| Acute Respiratory Distress Syndrome | Diagnosed by ICD-10-CM code J80. |
| Asthma | Diagnosed by an ICD-10-CM code beginning with J45. |
| Atrial Fibrillation | Diagnosed by an ICD-10-CM code beginning with I48. |
| Cancer Flag | Diagnosed by an ICD-10-CM code beginning with C. |
| Cerebral Infarction | Diagnosed by an ICD-10-CM code beginning with I63. |
| Chronic Kidney Disease | Diagnosed by ICD-10-CM codes E08.22, E09.22, E10.22, E11.22, E13.22 or beginning with I12, I13, N18. |
| Chronic Viral Hepatitis | Diagnosed by an ICD-10-CM code beginning with B18. |
| Chronic Obstructive Pulmonary Disease | Diagnosed by ICD-10-CM codes beginning with J41, J43 or J44. |
| Coronary Artery Disease | Diagnosed by ICD-10-CM codes beginning with I21, I22, I23, I24 or I25. |
| Crohns Disease | Diagnosed by ICD-10-CM codes beginning with K50. |
| Diabetes | Diagnosed by ICD-10-CM codes beginning with E08, E09, E10, E11, E13, O24.0, O24.1, O24.3 or O24.8. |
| Heart Failure | Diagnosed by ICD-10-CM codes beginning with I50. |
| HIV Flag | Diagnosed by ICD-10-CM codes B20, B97.35, O98.7, O98.71, O98.711, O98.712, O98.713, O98.719, O98.72, O98.73 or Z21. |
| Hypertension | Diagnosed by ICD-10-CM code I10. |
| Intracerebral Hemorhage | Diagnosed by an ICD-10-CM code beginning with I61. |
| Obesity | Diagnosed by ICD-10-CM code E66.1, E66.2, E66.8 or E66.9. |
| Obstructive Sleep Apnea | Diagnosed by ICD-10-CM code G47.33. |
| Ulcerative Colitis | Diagnosed by an ICD-10-CM code beginning with K51. |

**Laboratory Tests (continuous variables measured from a patient's blood sample, unless a different sample source is specified)**

| **Feature** | **Description** | **Unit** |
|---|---|---|
| Albumin | Amount of albumin | gram/deciliter |
| Alanine Transaminase | Amount of alanine transaminase. | unit/liter |
| Anion Gap | A measure of acid-base balance | milliequivalent/liter |
| Aspartate Aminotransferase | Amount of aspartate aminotransferase | unit/liter |
| Basophil (Count) | Count of basophil in white blood cell | ($10^{-3}\times$ count)/microliter |
| Basophil (Percentage) | Percentage of basophil in white blood cell | percentage |
| Blood Urea Nitrogen | Amount of nitrogen in the waste product urea | milligram/deciliter. |
| C-reactive Protein | Amount of C-reactive protein | milligram/liter |
| Calcium | Amount of calcium | milligram/deciliter |
| Chloride | Amount of chloride | milliequivalent/liter |
| $CO_2$ Total | Amount of carbon dioxide | milliequivalent/liter |
| D-dimer | Fibrinogen equivalent units of d-dimer | microgram/milliliter |

(continued on the next page)

Table 1: (continued from the previous page)

| | | |
|---|---|---|
| Estimated Glomerular Filtration Rate | The estimated glomerular filtration rate | milligram/minute/ ($1.73\times$meter-squared) |
| Eosinophil (Count) | Count of eosinophils in white blood cells | ($10^{-3}\times$ count)/microliter |
| Eosinophil (Percentage) | Percentage of eosinophils in white blood cells | percentage |
| Ferritin | Amount of ferritin | nanogram/microliter |
| Glucose | Amount of glucose | milligram/deciliter |
| Venous $HCO_3$ | Amount of bicarbonate in venous blood | milliequivalent/liter |
| Hemoglobin | Amount of hemoglobin | gram/deciliter |
| Lactate Dehydrogenase | Amount of lactate dehydrogenase | unit/liter |
| Lymphocyte (Count) | Count of lymphocytes in white blood cells | ($10^{-3}\times$ count)/microliter |
| Lymphocyte (Percentage) | Percentage of lymphocytes in white blood cells | percentage |
| Mean Corpuscular Hemoglobin | Average amount of corpuscular hemoglobin | gram/deciliter |
| Mean Corpuscular Hemoglobin Concentration | The average concentration of corpuscular hemoglobin | gram/deciliter |
| Mean Corpuscular volume | Average corpuscular volume | femtolitre |
| Mean Platelet Volume | Average platelet volume | femtolitre |
| Monocyte (Count) | Count of monocytes in white blood cells | ($10^{-3}\times$ count)/microliter |
| Monocyte (Percentage) | Percentage of monocytes in white blood cells | percentage |
| Neutrophil (Count) | Count of neutrophils in white blood cells | ($10^{-3}\times$ count)/microliter |
| Neutrophil (Percentage) | Percentage of neutrophils in white blood cells | percentage |
| Venous $O_2$ Saturation | Oxygen saturation in venous blood | percentage |
| Venous $PCO_2$ | The partial pressure of carbon dioxide in venous blood | mmHg |
| Venous pH | The pH of venous blood | No unit |
| Platelet | Amount of platelets | ($10^{-3}\times$ count)/microliter |
| Venous $PO_2$ | The partial pressure of oxygen in venous blood | mmHg |
| Potassium | Amount of potassium | millimoles/liter |
| Procalcitionin | Amount of procalcitionin | nanogram/milliliter |
| RBC Count | Red blood cell count | ($10^{-6}\times$ count)/microliter |
| Serum Creatinine | Amount of creatinine | milligram/deciliter |
| Sodium | Amount of sodium | millimoles/liter |
| Total Bilirubin | Total amount of bilirubin | milligram/deciliter |
| Total Protein | Total amount of two classes of proteins (albumin and globulin) | gram/deciliter |
| Troponin I | Amount of troponin I | nanogram/milliliter |
| WBC | Amount of white blood cells | ($10^{3}\times$ count)/milliliter |
| **Vital Signs (maximum and/or minimum of continuous-valued measurements during a patient's hospital encounter)** | | |
| **Feature** | **Description** | Unit |
| Maximum Diastolic BP | Maximum diastolic blood pressure | mmHg |
| Minimum Diastolic BP | Minimum diastolic blood pressure | mmHg |

(continued on the next page)

Table 1: (continued from the previous page)

| | | |
|---|---|---|
| Maximum Heart Rate | Maximum heart rate | Beats/minute |
| Minimum Heart Rate | Minimum heart rate | Beats/minute |
| Minimum $O_2$ Saturation | Minimum oxygen saturation | Percentage |
| Maximum Respiratory Rate | Maximum respiratory rate | Breaths/minute |
| Maximum Systolic BP | Maximum systolic blood pressure | mmHg |
| Minimum Systolic BP | Minimum systolic blood pressure | mmHg |
| Maximum Temperature | Maximum temperature | Fahrenheit |

Table 2: Ten highest contribution features for predicting mortality due to COVID-19 identified using the XGBoost method in Vaid *et al.* (2020)'s study (details of the features are in Supplementary Table 1).

| **Modality** | **Feature** |
|---|---|
| Admission | Age |
| Laboratory Tests | Troponin I |
| Laboratory Tests | Platelet |
| Vital Signs | Minimum $O_2$ Saturation |
| Laboratory Tests | C-Reactive Protein |
| Laboratory Tests | Aspartate Aminotransferase |
| Laboratory Tests | Glucose |
| Laboratory Tests | Calcium |
| Laboratory Tests | Blood Urea Nitrogen |
| Laboratory Tests | Procalcitonin |

### References


Radivojac, P. *et al.* (2013). A large-scale evaluation of computational protein function prediction. *Nature Methods*, **10**(3), 221–227.

Vaid, A. *et al.* (2020). Machine learning to predict mortality and critical events in a cohort of patients with covid-19 in new york city: Model development and validation. *J Med Internet Res*, **22**(11), e24018.

Walsh, I. *et al.* (2021). DOME: recommendations for supervised machine learning validation in biology. *Nature methods*, **18**(10), 1122–1127.

Whalen, S. *et al.* (2016). Predicting protein function and other biomedical characteristics with heterogeneous ensembles. *Methods*, **93**, 92–102.
